# A massively multi-scale approach to characterising tissue architecture by synchrotron micro-CT applied to the human placenta

**DOI:** 10.1101/2020.12.07.411462

**Authors:** W. M. Tun, G. Poologasundarampillai, H. Bischof, G. Nye, O. N. F. King, M. Basham, Y. Tokudome, R. M. Lewis, E. D. Johnstone, P. Brownbill, M. Darrow, I. L. Chernyavsky

**Author notes:** These authors contributed equally to the work (W. M. Tun, G. Poologasundarampillai). Joint lead authors (P. Brownbill, M. Darrow, I. L. Chernyavsky).

## Abstract

Multi-scale structural assessment of biological soft tissue is challenging but essential to gain insight into structure-function relationships of tissue/organ. Using the human placenta as an example, this study brings together sophisticated sample preparation protocols, advanced imaging, and robust, validated machine-learning segmentation techniques to provide the first massively multi-scale and multi-domain information that enables detailed morphological and functional analyses of both maternal and fetal placental domains. Finally, we quantify the scale-dependent error in morphological metrics of heterogeneous placental tissue, estimating the minimal tissue scale needed in extracting meaningful biological data. The developed protocol is beneficial for high-throughput investigation of structure-function relationships in both normal and diseased placentas, allowing us to optimise therapeutic approaches for pathological pregnancies. In addition, the methodology presented is applicable in characterisation of tissue architecture and physiological behaviours of other complex organs with similarity to the placenta, where an exchange barrier possesses circulating vascular and avascular fluid spaces.

## I. INTRODUCTION

Understanding complex vascular-rich organs such as the placenta has traditionally necessitated the adoption of multiple parallel imaging-based approaches, applied correlatively to gain structural and functional understanding of the tissue. These approaches generally require balancing resolution with sample volume. Structural analyses of placental vessels from whole organ or single villous branches have been conducted using multiple imaging modalities [1, 2] including CT angiography [3], micro-CT [4, 5], confocal laser scanning microscopy [6] and fluorescent CLSM [7]. Each of these methods has positives and negatives associated with it, however, none of them provide access to 3D micro-structures *in situ*, at high enough resolution to resolve the blood vessels, or functional units of the placenta, and still provide contextual information of the tissue.

Segmenting biological image data is a major challenge due to the structural complexity and intra/inter sample heterogeneity [8, 9]. Manual segmentation is commonly used for biological data segmentation, however, as the number and size of datasets increases this approach has become increasingly impractical [10]. Thus, an automatic or semi-automatic segmentation algorithm with high accuracy is required for high throughput segmentation of biological structure.

As one of the most complex vascular organs of the human body, the placenta is a well-suited model system for development of 3D imaging pipelines. The human placenta is an exchange organ with a large surface area of the feto-maternal interface packed in a relatively small volume, and an extensive feto-placental vascular network [11]. Its tightly integrated structural constituents span the spatial range from ∼10^−6^–10^−1^ m, necessitating a truly massive multi-scale imaging modality. Thus, new experimental and theoretical approaches are needed to bridge the microstructure of the placental exchange barrier and its macroscopic organ-level function, including the three-dimensional characterisation of the mesoscopic (between ∼0.1–1 mm) tissue domain [1, 11].

In this study, synchrotron X-ray imaging is used in combination with various sample processing conditions, including tissue contrast agents, vascular cast resins, fixation and embedding methods to generate high resolution massively multi-scale datasets of the human placenta. We then apply machine learning-based segmentation techniques for robust and efficient decomposition of maternal and fetal micro-domains in the large (≈8 mm^3^) datasets. Finally, spatial statistics and flow simulations of the feto-placental vascular network and associated intervillous porous space of the placental tissue are presented, and the results are validated against other modalities such as traditional 2D stereology analysis and *in vivo* magnetic resonance imaging.

The developed protocols for 3D multi-domain characterisation of tissues presented here, using the human placenta as an example, will enable more direct hypothesis-testing of the structure–function relationship in other organs where there are complex physiological fluidic/exchanger systems, such as in the kidney, lung, lymphatics, spleen, central nervous system, gut, bone-marrow and in wound healing and tumour biology.

## II. RESULTS

### A. Morphological study of mesoscopic placental tissue

#### (i). Comparative analysis of tissue preparation for X-ray micro-tomography (micro-CT) of complex soft tissues

Preparation of placental specimens for synchrotron micro-CT requires careful fixation, perfusion, staining and dehydration/embedding (Figure S1). Here we apply and qualitatively evaluate various specimen preparation methods to successfully image the complex architecture of human placenta (Figure 1). Tissue zinc-based fixative Z7 applied to all specimens provides tissue contrast when in-line phase contrast synchrotron imaging is performed (Figure S2). Specimen 2 (Figure S2A, D, G-J), fresh frozen by plunging in liquid nitrogen, preserves the physiological structures. Placental architectures, including a well resolved syncytiotrophoblast, blood vessels, capillaries, red blood cells and stroma can be differentiated in the 2D cross sectional and 3D rendered images. The syncytiotrophoblast appears as a thin (∼3 µm) envelope around an intermediate villous (Figure 2G). It is also possible to observe aggregated nuclei within a syncytial knot (Figure 2G-J). In Specimen 3 (Figure S2B, E, K-L) which was fixed with tissue fixative Zinc-7, ethanol dehydrated and wax-embedded, blood vessels, stroma and separately resolved microvillous and syncytiotrophoblasts are visible. Rendered 3D volumes in Figure S2K seem to show the presence of pores/open channels on the syncytiotrophoblast [12] which envelopes an intermediate villous circled in Figure S2L. From here, a machine-learning algorithm (U-Net) can be applied to segment and quantify the intervillous space (IVS) but not the vascular network or the stroma. Additional staining with 1% phosphotungstic acid (PTA) solution can further enhance the contrast for stroma (Figure S2F; Specimen 4) making it segmentable. PTA solution has low viscosity and so infiltrates into the IVS but takes several days (3 days for an 8 mm3 sample) to provide good contrast and signal-to-noise.

**FIG. 1:**
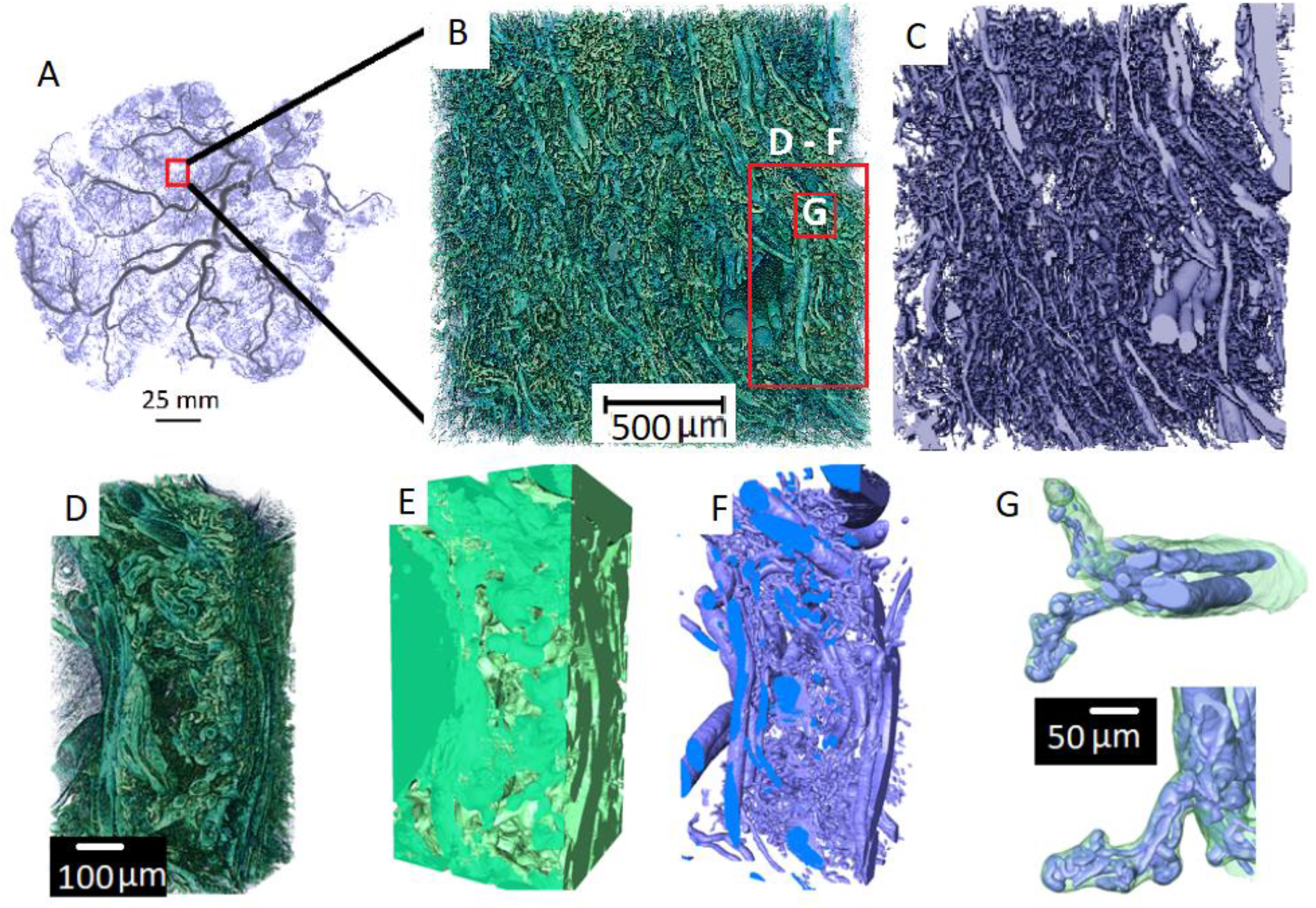
Multi-scale tissue architecture of placental tissue from synchrotron micro-CT. (**A**) Placental cast under micro-CT [unpublished image from [5]]. (**B-G**) Images demonstrating the complex hierarchical architecture of the human placenta (Specimen 1, normal placenta at term). (**B**) 3D rendering of ≈8 mm^3^ human placental tissue. (**C**) Fetal vascular network segmented from placental tissue using a U-Net algorithm. (**D-F**) A small section of the placental tissue was cropped from the original dataset (red box in *B*) and 3D rendered. (**D**) 3D rendering of ≈0.2 mm^3^ tissue showing different hierarchical features. (**E**) U-Net segmented fetal tissue component, (**F**) fetal vessels and (**G**) fetal capillary network with surrounding villous tissue overlaid.

**FIG. 2:**
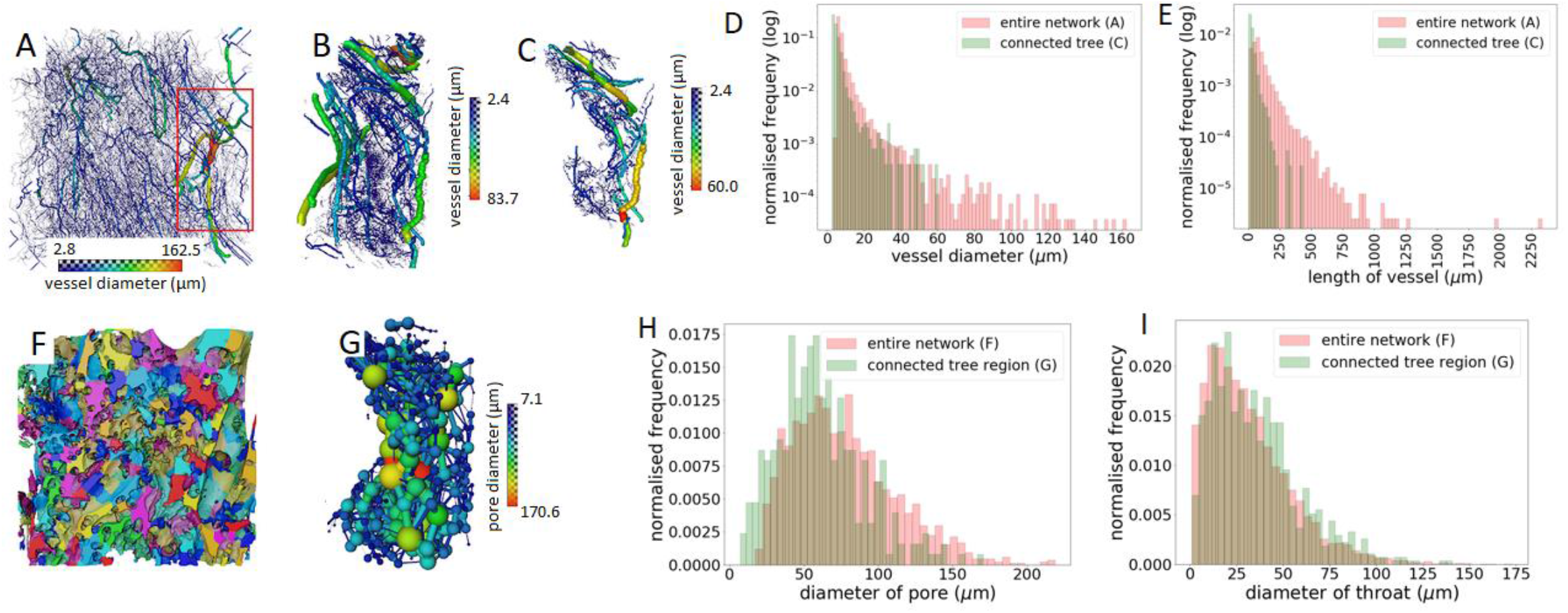
3D analysis of the structure of the fetal vascular network and maternal porous space (Specimen 1, normal placenta at term). (**A & B**) Skeletonized vascular structure from the entire tissue volume and from a small cropped region (the red box in *A*). (**C**) shows a single connected vascular tree in the cropped region shown in *B*. (**D & E**) Distributions of vessel diameter and length of blood vessels from the entire fetal vascular network and the connected vascular tree shown in *A* and *C* respectively. (**F**) Porous regions in the central tissue region (≈1.8 mm^3^). The colors represent different porous regions but are not related to the sizes of the pores. (**G**) Ball and stick model to represent pores and throats in the cropped region as in *C*. (**H & I**) Distributions of diameters of porous regions and connecting throats in the central tissue region and the region that encompasses the single connected tree in *C*.

In order to confidently segment the complex 3D vascular network resin infiltration casting via the fetal villi network is required to give sufficient contrast and phase differences. Specimens 1 (Figure S2C) and 3 (Figure S2B, E, K-L) were Zinc-7 fixed and perfused via the fetal side with casting resins (Batson’s and Yasuaki’s resins for Specimen 1 and 3, respectively), then ethanol dehydrated and wax-embedded. Batson’s contrast agent penetrates deeply into the fetoplacental circulation (Figure S2C) but shrinks inside larger vessels and possibly over-inflates smaller vessels, due to the high exertion force needed to infuse the resin via the fetal arterial cannula. Yasuaki’s resin is less viscous and highly has high X-ray attenuating than Batson’s and shrinks less, thereby likely better preserving vessel diameters. Both agents give good contrast to highlight the fetal lumen (Figure S2B, C). The viscosity of both Batson’s and Yasuaki’s reagents make them unfavourable for infusion into the IVS since filling is incomplete. Extent of tissue processing influences final architecture. Ethanol dehydration and wax embedding (Specimen 3 and 4) may introduce tissue deformation, in particular the villous trophoblast appears to delaminate from the villi stroma (Figure S2B, E). Tissue deformation occurred to much less extent with CP-dried specimens (Specimen 1; Figure S2C). Whilst, cryo-freezing seems to have caused syncytiotrophoblast shedding to occur, but overall it has preserved the morphology of specimens more than the other methods. However, synchrotron micro-CT imaging of cryo-frozen specimens requires a cryo-stream or cold stage to maintain the specimen at cryo-condition during scanning. Figure S1A provides a summary of various sample processing conditions tested here and a flow chart (Figure S1B) that recommends appropriate processing pipelines to resolve the multiscale structures in the placenta.

#### (ii). Characterisation of feto-placental vascular network in 3D

Feto-placental vascular network and porous materno-placental tissue domain were quantified in 3D for Specimen 1 (normal) and Specimen 2 (pregnancy with fetal growth restriction, FGR). The scanned 9 mm3 cubed samples generated ≈ 8 mm^3^ of digital 3D data. The entire volume (14.2 billion voxels) was segmented and analysed to identify placental tissue architecture across multiple scales (Figure 1 and Supplementary Video 1). A disambiguated 3D rendering (Figure 1B) from the entire volume of tissue demonstrated the sample tissue volume and three-dimensionality achieved with the technique, while the detail in the terminal capillary loops and surrounding villous tissue (Figure 1G) demonstrated the resolution. Two separate 2D U-Nets (Figure S1) were used alongside expert-generated training data (Figure S3) to fully segment the feto-placental vascular network and the maternal and fetal blood volumes (Figure 1C, E, F, G).

Once the complex tissue was segmented, multiple data analysis pipelines and simulations could be run, across the imaging scales. Figure 2 displays the skeletonized vascular structure from the entire tissue dataset (2A), a small cropped region (2B) and a single connected tree (2C). The median (interquartile range; IQR) diameter of blood vessels in fixed tissue is 6.8 (IQR: 6.2–8.6) µm for the entire network and 4.4 (IQR: 3.4–6.0) μm for a single connected tree (Figure 2D), while the length of blood vessels is 73.8 (IQR: 47.4–115.6) μm for the entire network and 21.2 (IQR: 13.6–34.1) μm for a single connected tree (Figure 2E). Tortuosity of the blood vessels is 1.2 (IQR: 1.1–1.4) for the entire network (Figure S4) and 1.2 (IQR: 1.1– 1.3) for a single connected tree. A limitation in the current analysis is that skeletonization and quantification of blood vessels for the entire network (Figure 2A) was carried out on down-sampled image data due to computational limitations. Therefore, the measurements from the entire vascular network are likely to be less accurate in comparison to those from the single connected tree, where the full resolution data was used.

### B. Characterisation of the porous materno-placental tissue domain in 3D

To characterize the maternal blood porous space, the centre region of the scan, ≈1.8 mm^3^from the entire data set (see supplementary Figure S3A) and the porous space that encompasses the single connected tree (from Figure 2C) were used. The median (interquartile range; IQR) diameter of pores is 72.2 (IQR: 50.6–97.2) μm for the central region and 57.6 (IQR: 40.6–77.8) μm for the single tree region. The diameter of the throat (a throat is a region that connects two individual pores) is 27.0 (IQR: 14.4–43.6) μm for the central region and 31.0 (IQR: 16.8–47.0) μm for the single tree region. The distributions of diameter of porous regions and connecting throats of both the central region and the region that surrounds the single connected tree (both from specimen 1) are shown in Figure 2H and 2I. The median and interquartile ranges of diameter of pore and throat, length of throat and number of connected pores were also analysed for both Specimens 1 and 2, and the results are presented in supplementary Figure S5. These analyses were performed on central tissue regions only.

The flow tortuosity of porous regions was plotted with different minimal lengths. The smallest minimal length employed here (85 μm) is bigger than the mean diameter of a pore (≈80 μm) since the blood flow inside a single pore is considered straight. Supplementary Figure S6G shows that the porous tortuosity falls between 1 and 3 in both Specimen 1 and 2.

### C. Mesoscopic flow analysis in the human placenta

Using Specimen 1, the maternal flow velocity in the IVS was simulated for a fixed pressure gradient applied in three principal directions (Figure 3 and Supplementary Video 2). The relationship between fetal tissue (Figure 3A(i)) and maternal flow (Figure 3A(ii)) in velocity map and streamlines respectively and the inter-relationship between maternal flow streamlines (Figure 3B) and fetal vascular network (Figure 3C) were visualized. The distribution of flow velocities (Figure 3D) were in the range 0–1670 (mean: 8) μm/s in the x direction, 0–1670 (mean: 10) μm/s in the y direction, and 0–1660 (mean: 7) μm/s in the z direction respectively.

**FIG. 3:**
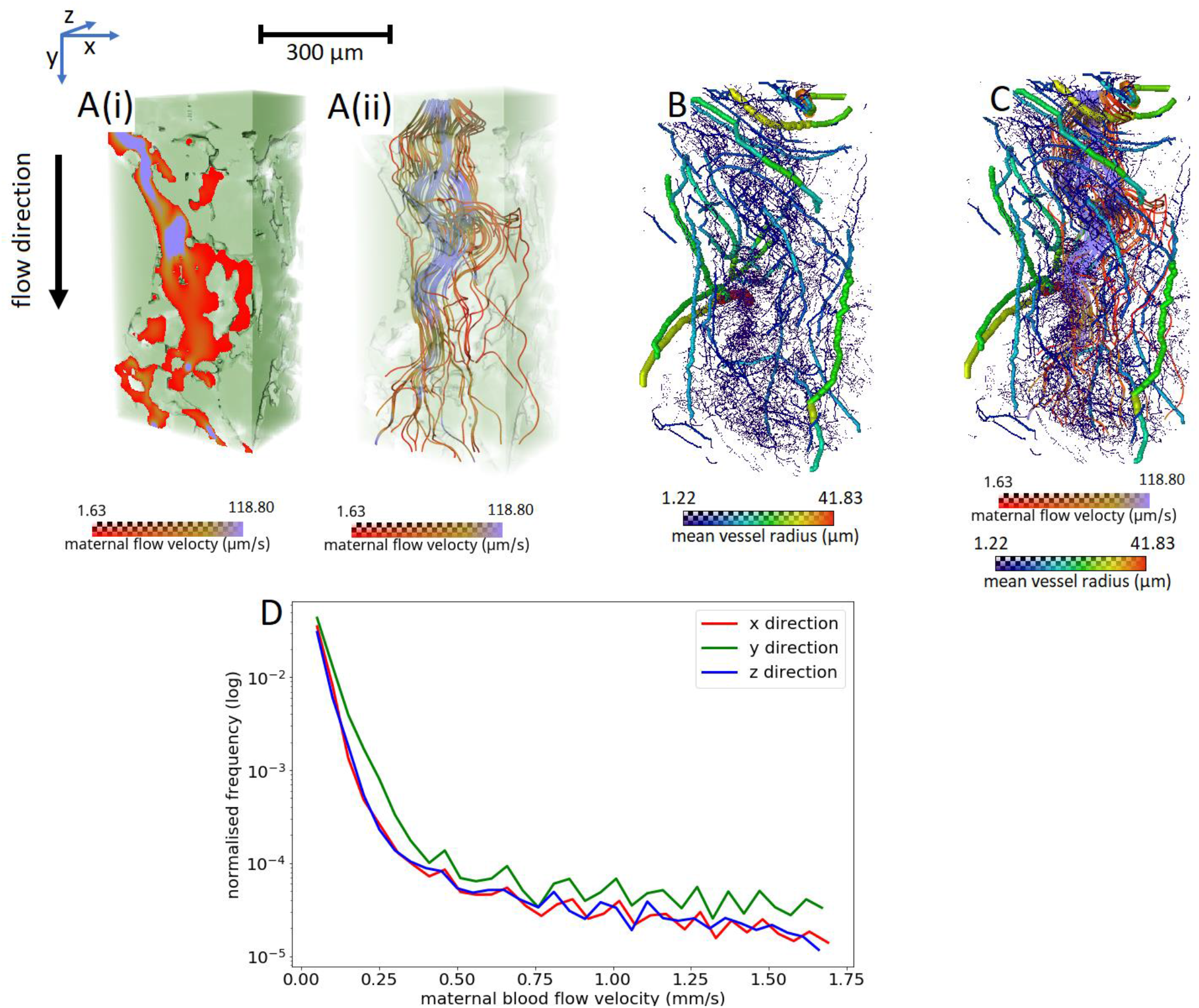
Flow simulations across maternal IVS to visualize the inter-relationship between maternal blood flow and fetal vascular flow (Specimen 1, normal placenta at term). (**A**) shows the maternal (i) flow velocity map and (ii) flow streamlines in the IVS. (**B & C**) show the inter-relationship of maternal flow streamlines and fetal blood vascular network. (**D**) Line graphs comparing the maternal blood flow velocity distributions in three different directions with a fixed pressure gradient. x-axis of the graph shows maternal blood flow velocity and y-axis shows the normalised frequency in logarithmic values.

Despite the approximately 1:2:1 (x:y:z) aspect ratio of the studied porous domain, the flow resistance in the y direction was found to be approximately twice as small as in the x and z directions, with the corresponding diagonal components of the empirical hydraulic permeability tensor ≈(1.5, 3.4, 2.0) μm^2^, indicating a relatively strong flow anisotropy of the IVS.

The connectivity of the IVS domain was also explored as a function of the minimal pore-throat diameter needed to connect a random central pore to the periphery of the domain (Figure S6A-F). Such critical diameter was found to be just about 25% of the pore size, pointing to a highly connected IVS that facilitates flow-limited transport at the tissue mesoscale of ∼1 mm (see S8 in the Supplementary Text for more details).

### D. Uncertainty quantification and scale-dependence of morphological metrics

Placental tissue area fraction fluctuates across the 3D volume in both specimens. Based on the central tissue region (≈1.8 mm^3^), the area fraction of Specimen 1 ranges from 0.54–0.73 (mean: 0.64) and that of Specimen 2 ranges from 0.58–0.71 (mean: 0.65; see Figure 4A and Supplementary Figure S7A, B). The tissue volume fractions for Specimens 1 and 2 are 0.64 and 0.65 respectively. Figure 4B shows how the standard deviation of the volume fraction decreases with increasing ROI size (see also Figure S7C). The scale-dependent error in fetal tissue volume fraction, specific surface area (Figure 4C) and the 2-point correlation function vs. distance for both specimens (Figure 4D) are also presented.

**FIG. 4:**
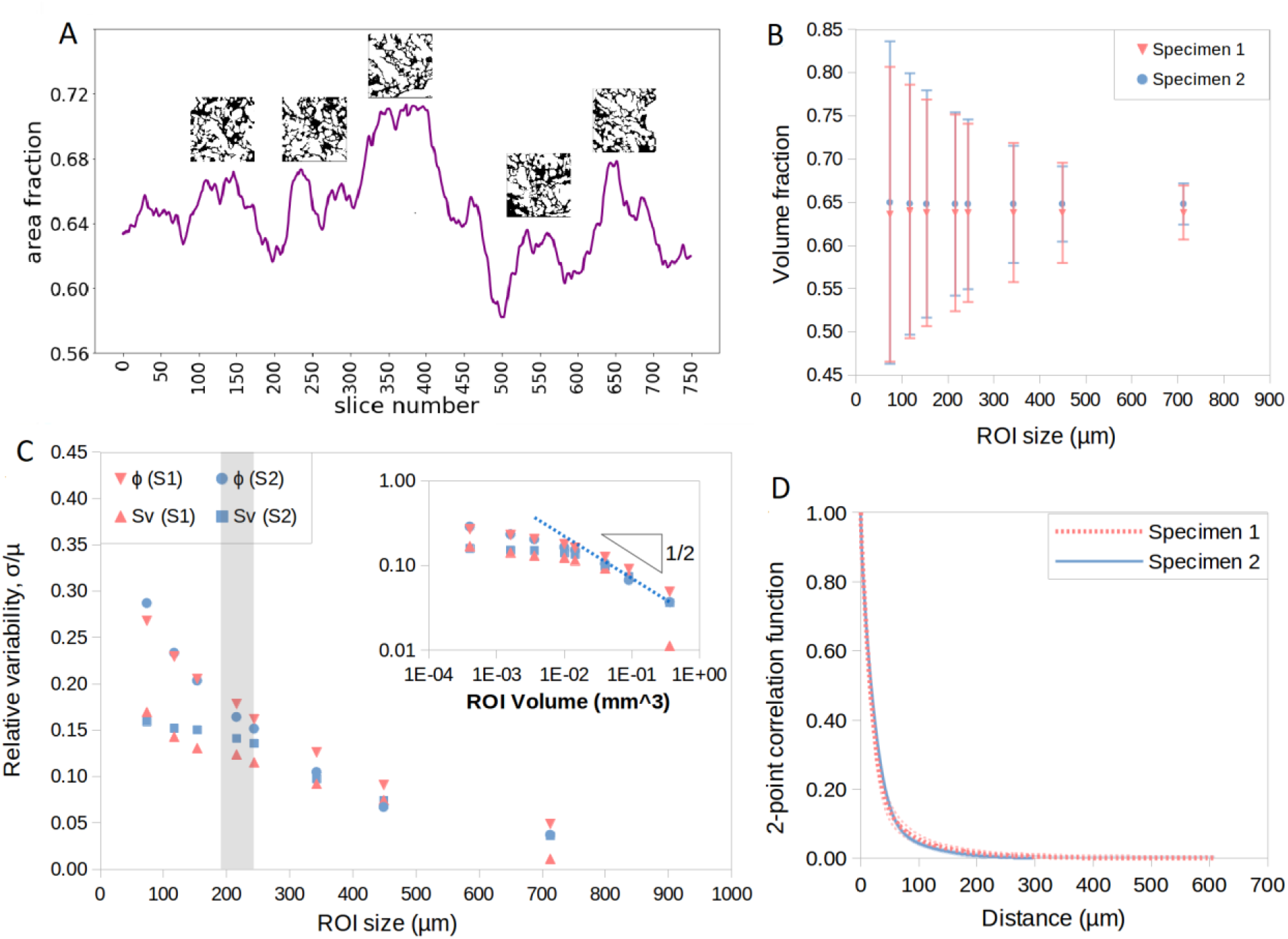
Uncertainty quantification and scale-dependence of morphological metrics (Specimens 1 & 2). (**A**) Fluctuations of placental villous tissue area fraction in Specimen 2 for the region of interest (ROI) with the volume of ≈1.22×1.22×1.22 mm^3^ (insets illustrate selected slices). (**B**) Volume fraction fluctuations (mean±SD) vs. effective ROI size. (**C**) Scale-dependence of a relative error in the tissue volume fraction (ϕ) and specific surface area (S_v_) estimates; the inset compares to the theoretical prediction of ∼(ROI Volume)^1/2^ on a logarithmic scale (see Supplement S9). (**D**) Radial two-point autocorrelation function (mean; SD is shown with shaded lines); the corresponding approximate transition range to uncorrelated meso-scale (mean autocorrelation ≲1%) is shown as a grey stripe in *C*.

Figure 4C illustrates that the relative magnitude of fluctuations in both the tissue volume fraction and specific surface area fall approximately inversely proportional to the square root of the ROI volume (σ/μ ∼V_ROI_^−1/2^) for the sufficiently large ROI size (compared to the characteristic lengthscale λ ∼200 μm; see Figure 4D and Table 1). This is in agreement with theoretical predictions for a generic porous medium (see S9 in the Supplementary Text for more details). Thus, to reduce the estimation error in morphological metrics by a factor of 2, a four-fold increase in the ROI volume is required. For smaller ROI volumes (i.e. when the ROI size is less than ca. 200 μm), the morphometric fluctuations are even more sensitive to the measurement scale.

**TABLE 1:**
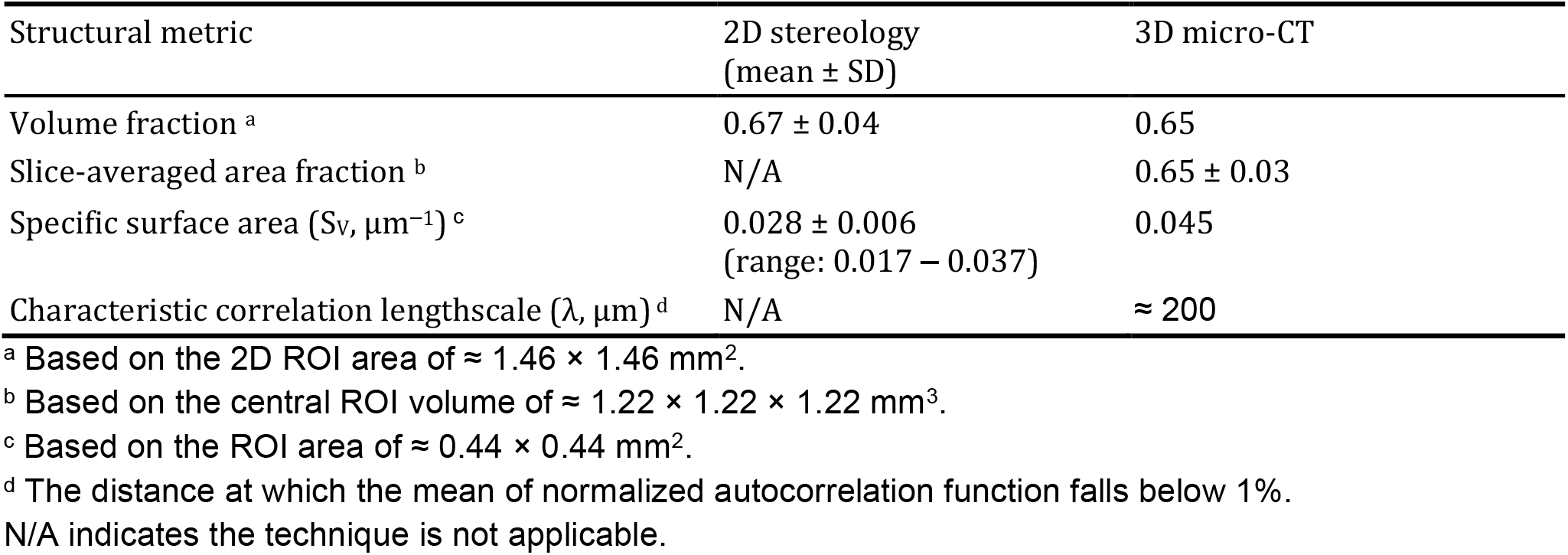
Comparison of 2D- and 3D-based placental tissue morphometrics (Specimen 2; see Figs S7 & S8 for more details).

Three-dimensional (3D) morphometrics were also compared to the estimates based on two-dimensional (2D) ‘virtual’ stereology (Figure S8) on the same dataset to cross-validate the estimated tissue volume fractions and specific surface areas between these two approaches. Table 1 reports the villous tissue volume fraction and specific surface area for Specimen 2 estimated by traditional 2D stereology and 3D morphological analysis. Comparison to the direct 3D estimates (Table 1) shows good agreement for the volume fraction but an up to 40% relative difference for the specific surface area.

## III. DISCUSSION

In this study, we present a novel approach to quantitatively characterise the 3D structure of complex mesoscale tissues using the human placenta as a model. We demonstrate how up to four orders of magnitude can be bridged in a single imaging dataset using synchrotron X-ray micro-tomography, and propose an optimized pipeline for tissue sample processing and image segmentation to robustly evaluate morphological data of complex vascular systems. To our knowledge, this is the first study to characterise the maternal placental IVS porous region and to demonstrate the inter-relationship between maternal blood flow and fetal vascular network. Finally, we compare traditional stereological approaches with direct 3D structural analysis and quantify the scale-dependent fluctuations in common morphological metrics. The developed framework helps to identify a minimal characteristic size for heterogeneous placental and other complex soft tissues and provides the tools to control the associated uncertainties for robust and scalable predictions.

The placental specimens in this study were prepared using different tissue processing techniques (critical point (CP)-drying, fresh-frozen, wax-embedding) and perfused with different contrast agents (Batson’s and Yasuaki’s; Table S1). CP-dried and wax-embedded specimens showed more deformation and shrinkage of villous tissue in comparison to fresh-frozen specimens that maintained the tissue at near-native anatomical condition. Among the three processing techniques, CP-dried specimens displayed the best image contrast between fetal tissue and maternal IVS (Figure S2). Yasuaki’s gave a superior contrast of the fetal vascular network than Batson’s, meaning Yasuaki’s-perfused specimens could be easily segmented with contrast-based image segmentation methods while Batson’s-perfused specimens require more complex segmentation strategies. However, one drawback with Yasuaki’s resin was lower perfusion efficiency which manifests as smaller arterioles and venules in the terminal capillaries. Sample processing must be considered in light of both the preservation of the region of interest, but also the downstream data analysis and segmentation strategy (Figure S1A). In an ideal case (Figure S1B, specimens would immediately be Z7 fixed, fetal network infiltrated with a resin, paraformaldehyde (PFA) fixed, and then cryo-frozen and micro-CT imaged under cryo-conditions. If a cryo-stage is not available then specimens should be ethanol dehydrated and CP-dried before imaging under ambient conditions. In order to segment the various structures a machine-learning algorithm (U-Net) should be employed. This proposed sample preparation and imaging pipeline would be applicable to other vascular-rich organs [13-15].

In recent years, automatic or semi-automatic segmentation methods using deep learning algorithms have become popular in the vascular biology [16] and placental biology fields [17]. Convolution neural networks (e.g. U-Net) have been successfully applied in segmenting various vascular systems including retina [18, 19], brain [20], whole placental volume [21], and larger placental vessels [22]. Here for the first time, we apply a U-Net towards multi-domain segmentation of the intricate fetal vascular network and maternal porous space, including terminal capillary loops where vessel diameters are smaller than 50 μm. The semi-automatic algorithm used here requires a minimal amount of training data (≈1% of entire volume). The most striking aspect of the U-Net segmentation is the detection of the smallest terminal capillaries, with diameters of the fixed tissue as small as ≈2.5 μm (IQR: 6.2–8.6). These tiny terminal capillaries are exceedingly difficult to segment manually, so their presence in the U-Net segmentation provides a less time-consuming and more consistent method for their inclusion in analyses.

The U-Net predictions were assessed by dice score coefficient, a commonly used voxel-based validation metric for image segmentation algorithms [23]. Using the dice score, comparisons between multiple manual expert segmentations of complex, biological components have varied between ≈0.27 and ≈0.69 [24]. When comparing the U-Net segmentations to expert manual validation segmentations, the dice scores for vascular and tissue regions are 0.8 and

0.9 respectively (Figure S9). There are, however, some artefacts present in the U-Net segmentation (mislabelling of maternal porous region as fetal tissue) that could lead to overestimation of the surface area of fetal tissue. In addition, the current U-Net model was not able to generalize across datasets, instead requiring manual training data from each individual dataset. This may be due to the variation in specimen processing and imaging, which manifested significantly different appearances in the final datasets.

Understanding the structural development of the placenta is critical since the common pregnancy disorders FGR and pre-eclampsia are associated with impaired development of the placental vascular network [25, 26]. The human placenta is hemochorial and possesses a dual blood supply; maternal and fetal circulations are separated, but work collaboratively as a multi-villous exchange system [27]. Defects in either supply could impair nutrient and gaseous exchange. Here, we have quantified the continuous fetal vascular networks from a ≈8 mm^3^ placental tissue block capturing the transition from the micro- to meso-scale, with the size range of blood vessels and intervillous pores of the order of 10–1000 µm (Figure 2). In order to appreciate the spatial inter-relationship between these two circulations, we have visualized the maternal flow intertwined with the fetal vascular network (Figure 3C). We simulated maternal blood flow in a large portion of the segmented intervillous pore space for a fixed pressure gradient of ∼10 Pa/mm within placental tissue [28, 29]. The predicted flow had an exponential distribution of velocities, typical for a random porous medium [30], with the range of ≈0–2 mm/s. This is consistent with previous MRI *in vivo* estimations that reported regions of slow (< 0.5 mm/s) and fast (> 1 mm/s) blood flow [31]. Furthermore, our results suggest a new hypothesis that local IVS pore sizes influence nutrient and gaseous exchange efficacy. Future studies are needed to identify whether IVS porosity spatially correlates with the calibres and orientation of associated villous microvessels, and to determine the extent of maternal-fetal placental flow matching. The impact of villous tissue compliance on flow distribution in the IVS, particularly, in the proximity of decidual arterial inflows, also warrant further investigation.

With the intention of understanding the normal physiological blood flow and deviations occurring in diseased pregnancies, numerous mathematical and computational models have been developed and attempt to replicate the utero-placental circulations [11, 32-34]. A particular challenge is the geometric complexity of intervillous maternal porous regions which are difficult to characterize [33]; therefore, there is a lack of knowledge on placental porous structure. Porosity within human tissues has been extensively investigated in bone and other soft tissues (e.g. adipose tissue) from the perspective of tissue engineering [35-38]. Nonetheless, the placental maternal IVS porous medium has been less well studied. Here, we have quantified the size of porous regions, the connecting throats between individual pores, and the length of connecting throats. Moreover, we report the number of coordination for individual pores, and also visually examine the connectivity by thresholding the size of connected regions (Figure S6). Both qualitative and quantitative analyses suggest that the maternal porous space is highly connected and there is no unconnected, isolated porous region. The minimal diameter of throat is 0.86 μm (IQR: 14.4–43.6), in line with the average thickness of a human red blood cell (0.8–1 μm), indicating blood flow could occur throughout the porous medium with some impedance [39]. However, the resistance to IVS flow would vary depending on the length and diameter of connecting throats/pores, suggesting a potential explanation for the heterogeneous maternal flow velocity reported in [29] and also in our flow simulations.

Contrast imaging and mesoscopic flow analysis reported here could readily be applied to other organs where it is challenging to understand structure-function relationships when a fluid circulates around epithelia or where flow is not confined to a vascularised system. This type of anatomy can be found in the intestine and the central canal of the spine. In the gut, uptake of nutrients and therapeutic compounds occur from the crypts of the jejunum lumen, passing into the systemic circulation having traversed associated villi. The crypt lumen has been observed to become vacuolated following chemotherapy, altering their porosity, which is hypothesised to affect the efficacy of nutrient and therapeutic uptake [40]. In a second example, the motile cilia of the spinal cord central column and brain generate an efficiency of cerebrospinal fluid flow that is sensed and regulates structural modelling of the shape of an organism during embryonic development [41]. Thus, the workflow we have developed could provide anatomical insights on tissue structure and function in homeostasis and pathology.

It is important to note that the pore diameters measured here could be influenced by the flow and the hydrostatic pressure within the IVS during perfusion fixing, compared to non-perfusion fixed tissue, where villous and IVS deflation could occur [42]. Following partum, the placenta collapses and loses a significant volume of maternal blood from the IVS. However, a more physiological pressure and flow within the IVS *in vivo* can be recapitulated during *ex vivo* dual perfusion of the human placenta, yielding a more normal spatial arrangement of villi compared to the arrangement produced by the direct immersion fixing of deflated placental material. Consequently, most historic stereological analyses have likely under-estimated the volume fractions of the IVS in this tissue [42]. In turn this will have led to an under-representation of pore size diameter and connectivity characteristics. However, potential confounders of the *ex vivo* perfusion model relate to an artificial rheology, created by a blood-free environment, with some compromise in viscosity linked to altered shear stress within the two circulatory systems. One unresolved question is how to determine the tissue sample size required to accurately obtain statistically-representative data. This is particularly true for placental tissue due to its inherent heterogeneous nature. While it is true that the larger the tissue section, the more information one can retrieve, using a large-sized sample comes with technical and economic burdens. Therefore, systematically characterising the minimal tissue size that could provide adequate information with the least error is advantageous. Using uncertainty quantification and scale-dependence of morphological metrics applied to a variety of placental tissue samples, we identified a transition from microscopic fluctuations to tissue-scale properties at the mesoscale of ∼200 μm (Table 1). Thus, the smallest ROI volume required, without failing to comprehend the heterogeneous nature of the finest features of exchange villi in placental tissue, is of the order of magnitude of 0.1 mm^3^ (see Figure 4). Although, this smallest cut-off point might vary in other tissues with different geometric properties (e.g. larger blood vessels), the generic scale-dependence of the magnitude of 3D morphometric fluctuations (inversely proportional to the square root of the ROI volume, extending the results of [43]) is shown to be sufficiently universal for a wide class of porous media found in biological solute and gas exchangers. Future work is needed to further refine and extend the developed framework to placental pathologies and to other organs.

In conclusion, this paper offers both a novel approach and a validated workflow for massively multi-scale characterization of soft tissues with complex vascular networks. In addition, several detailed morphology and functional characteristics of our chosen model tissue, placenta, have been analysed in 3D. In future, these methods could be used to explore underlying causes of disease and inform potential future treatments.

This approach was enabled by a combination of innovative sample preparation, advanced synchrotron-based imaging, and state-of-the-art segmentation algorithms that bridge the gap of previously disconnected characterisation strategies using traditional X-ray tomography and angiography (cf. Figure 1A), light microscopy (cf. Figure 1B) and confocal or electron-microscopy (cf. Figure 1F). In the placental research arena the data generated here represents, for the first time, a multi-scale approach applied to characterise the architecture of both maternal and fetal domains by synchrotron micro-CT. Additionally, the recommendations provided for future researchers should allow translating the workflow to quantify structures in other complex, soft tissues.

## IV. METHODS

### A. Specimen collection and preparation

#### *Ex vivo* placental perfusion and preparation of tissue blocks

*Ex vivo* dual perfusion of human placental cotyledons was conducted as previously described [42] from two pregnancies complicated by FGR and two from uncomplicated pregnancies (see Table S1). Briefly, placentas were collected within 30 minutes of delivery and the chorionic plate artery and vein corresponding to an intact cotyledon were each cannulated. A suitable cotyledon was selected on the basis of intactness following delivery. Peripheral cotyledons were not excluded. Fetal-side perfusion was commenced in an open circuit at 6 mL/min with a modified Earle’s bicarbonate buffer, gassed with 5% CO_2_/bal. N_2_. Following a quality control check at T=5 minutes that fetal-side venous outflow was ≥ 75% of fetal-side inflow, the maternal-side perfusion was commenced in open circuit at 14 mL/min with the same buffer, gassed with 21% O_2_/5% CO_2_/bal. N_2_, via a single glass cannula (i.d. =2 mm) held 5 mm below the decidua surface. The maternal cannula was inserted at the centre of a placental cotyledon. For the tissue mass of ca. 30-40 g, this typically represented a random location of the insertion with respect to several villous trees within a cotyledon, below the decidual surface. The cotyledons continued to be perfused in open circuit for three hours prior to perfusion fixation from the maternal surface with Zinc 7 fixative [44] for 15 minutes. Various contrast reagents were infused into the fetal or maternal circulatory compartments (Table S1).

Colloidal dispersion of NiAl layered double hydroxide (LDH) “Yasuaki” resin was prepared following our previous report [45]. Briefly, NiCl_2_·6H_2_O and AlCl_3_·6H_2_O were dissolved in a mixture of EtOH and ultra-pure water followed by the addition of acetylacetone (acac). To this mixture, propylene oxide (PO) was added as an alkalization agent [46] and the container was sealed and kept at a room temperature (RT ∼20°C). The obtained suspension was kept in a freezer (−20°C) and then dried under a vacuumed condition (<10 Pa), yielding dried NiAl LDH nanoparticles. To make up 5 mL of resin, the powdery NiAl LDH (1.0 g) was dispersed in ethanol (EtOH; 2.5 mL), and then methyltriethoxysilane (MTES; 2.0 mL) and tetraethyl orthosilicate (TEOS; 0.5 mL) were added to this mixture. Then, H_2_O (0.7 mL) was added just before perfusion to initiate the gelation reaction. The procedures were performed under stirring at RT. This resin was applied immediately after fixing the lobule *in situ* within the perfusion cabinet, ahead of any other contrast reagents that might have been used.

On occasions a Batson’s resin from the “Batson’s no. 17 Anatomical Corrosion Kit” (Polysciences, Inc., Europe) was applied to the fetal circulation following perfusion. The liquid resin was prepared as a 20 mL base solution, 2.4 mL of catalyst and one drop of promoter; and manually injected via the chorionic plate arterial cannula until emergent at the chorionic plate vein cannula. Both cannulae were clamped and the whole lobule was allowed to polymerise overnight on iced water, within a sealed plastic bag.

The postperfusion fixed and post-contrast-infused cotyledon was excised from the non-perfused tissue and a 5 mm vertical slice of the cotyledon, adjacent to the inflow locus of the maternal cannula, was dissected from the cotyledon and fixed in a PFA fixative overnight to stiffen and further preserve the tissue. In all cases, following PFA diffusion fixing, 5 mm sized cubes were dissected from the vertical tissue sections and stored in sterile PBS. In some cases, the small blocks were infused with a further contrast agent, phosphotungstic acid (PTA), for several days or hours. In other cases, the cubes underwent CP-drying using an E3100 Critical Point Dryer (Quorum, UK), following manufacturer’s instructions, and dipped into liquid nitrogen to freeze fracture a 3 x 3 mm^2^ cross-sectional sample for imaging, or prior to wax embedding.

The methods for processing whole-organ vascular casts (see Figure 1A) were as previously described [5].

#### 3D tomographic imaging

At the Diamond Light Source (DLS) facility (Harwell, UK; Manchester Imaging Branchline, I13-2), in-line high-resolution synchrotron-sourced phase contrast micro-computed X-ray tomography was used to generate images, following various methodologies to optimise image quality and feature extraction [47, 48]. Briefly, micro-computed tomography employed filtered (1.3 mm pyrolytic graphite and 3.2 mm Al filters) polychromatic X-ray beams with energy in the range of 8−30 keV to probe the samples. Transmitted X-rays produce visible light on striking a scintillator (500 μm thick CdWO4) positioned between 60–100 mm away, in-line with the sample. The light was then magnified with various objectives and imaged on a sCMOS (2560×2160 px) detector (pco.edge 5.5; PCO AG, Germany). Optical magnification of 8x was employed resulting in an effective isotropic px size of 0.81 μm. In total, 3001−4001 X-ray projections were recorded over 0−180° rotation using exposure times between 80–200 ms. Projections were reconstructed into 3D datasets using a filtered-back projection algorithm [49] incorporating dark- and flat-field correction, ring artefact suppression and lens-blurring [50, 51].

Samples (3×3×3 mm3 cubes) were prepared following various methodologies (see above and Table S1) to optimise image quality and feature extraction. Specimen 1 was ethanol dehydrated then CP-dried and imaged at RT (60 mm S-D distance, 3001 projections, 0.08 s exposure). Specimen 2was plunge frozen into liquid nitrogen and imaged (60 mm S-D distance, 4001 projections, 0.15 s exposures) whilst maintaining a sample temperature of −20°C using a cold stage [52]. Specimen 3 was ethanol dehydrated then wax-embedded and imaged at RT (100 mm S-D distance, 4001 projections, 0.2 s exposure). Specimen 4 was PTA stained, ethanol dehydrated then wax-embedded and imaged at RT (10 mm S-D distance, 4001 projections, 0.2 s exposure).

#### B. Segmentation

‘Ground-truth’ training data for Specimen 1 was created using SuRVoS (Super Region Volume Segmentation Workbench) which applies a supervoxel segmentation strategy as described in [53]. A 256×256×256 px (0.21×0.21×0.21 mm) region of the full resolution volume was segmented for the feto-placental vascular network and a 384×384×384 px (0.31×0.31×0.31 mm) region was segmented into the maternal and fetal blood volumes. These segmented regions along with the corresponding image data were then used to train two separate 2D U-Net models for binary segmentation [54]. All aspects of model construction, training and data prediction were done using the fastai deep learning Python library [55]. The U-Net model architecture used a ResNet34 [56] encoder that accepts images of size 256×256 px and that had been pre-trained on the ImageNet dataset [57]. For model training, both the segmented label volumes and the corresponding data volumes were sliced into 2-dimensional images parallel to the xy, xz and yz planes. For each model, a randomised dataset was created from the pool of images with an 80%/20% split between training and validation image sets. The default fastai image transformations and augmentations were used. Model training was carried out using binary cross-entropy as the loss function and evaluated using the Intersection over Union (IoU/Jaccard) score as the metric on the validation set. For the vascular network data, training was carried out for 10 epochs giving a final IoU score of 0.93 on the validation set, the loss for the training set was 0.056 and the corresponding loss for the validation set was 0.052. For the maternal/fetal blood volumes, training was carried out for 15 epochs, giving a final IoU score of 0.93 on the validation set, the loss for the training set was 0.099 and the corresponding loss for the validation set was 0.097.

To overcome the issues of using a 2-dimensional network to predict 3-dimensional segmentation, a data-averaging approach was developed. To generate the vascular network and maternal/fetal blood volume segmentations for the full 2520×2520×2120 px (2.05×2.05×1.72 mm) data, the image data volume was sliced into three stacks of 2-dimensional images parallel to the xy, xz and yz planes. The corresponding segmentation for each of these image stacks was predicted before being recombined back into a 3-dimensional dataset, thereby producing 3 separate segmented volumes. The image data volume was then rotated by 90 degrees around the 4-fold symmetry axis running perpendicular to the xy plane and the entire slicing and prediction process was repeated again. After 4 cycles of this process, the resulting 12 segmented volumes were summed and a final segmentation produced by applying a threshold to the data where there was agreement between 6 and more of the predictions in the case of the maternal/fetal blood volumes and between 9 and more of the predictions in the case of the blood vessels.

To segment the fetal tissue components from Specimen 2, the central region of the full resolution dataset (2520×2520×2120 px = 2.05×2.05×1.72 mm) was cropped to obtain a volume of 1500×1500×1500 px (1.22×1.22×1.22 mm) (see Supplementary Figure S3A), which was then down-sampled to obtain a volume of 750×750×750 px. This down-sampled, cropped volume was then split into eight sub-volumes, each of which were manually segmented using SuRVoS (Supplementary Figure S3B to S3D).

Segmentation and analysis along with U-Net training and prediction were carried out on a High-Performance Computing Cluster node. The node had two Intel® Xeon® Gold 6242R Processors each with 20 cores running at 3.1 GHz, and 768 GiB of system memory. The GPU used was an NVIDIA Tesla V100 with 32 GB of available memory.

### C. Validation of semi-automated segmentation by U-Net

To validate the U-Net prediction of fetal vascular network in Specimen 1, two different regions of 384×384×384 px (0.31×0.31×0.31 mm), which were located away from the original training data, were randomly chosen and SuRVoS-segmented manually. The similarities between the SuRVoS-segmentations and the U-Net predictions (Supplementary Figure S9A and S9B) were compared using the Sørensen–Dice score = 2|M ∩ U|/(|M| + |U|), which compares the area of the overlap (M ∩ U) to the average area of the manually-segmented (M) and U-Net-predicted (U) regions respectively (estimated for each slice).

To validate the U-Net prediction of the fetal tissue component in Specimen 1, a similar strategy was used. Dice scores were again calculated between two SuRVoS-segmented Validation Regions (256×256×256 px; 0.21×0.21×0.21 mm) and the U-Net prediction (Supplementary Figure S9C and S9D). The fetal tissue area fraction of individual slices was also compared between U-Net predictions and SuRVoS segmentations.

Supplementary Figure S9E and S9F show the Dice scores for fetal vascular network segmentation and fetal tissue segmentation respectively. The mean dice score for vascular validation Region #1 is 0.81 and that for vascular validation Region #2 is 0.88. Vascular Region #2 is likely to have a higher dice score because it contains fewer but larger vascular branches in comparison to vascular Region #1. The mean Dice scores for tissue validation Region #1 and Region #2 are 0.97 and 0.96 respectively. A higher dice score in the tissue regions as opposed to the vascular regions is expected because the vascular network segmentation has more boundaries than the tissue segmentation, where most discrepancies between U-Net and SuRVoS segmentation occur. The area fraction comparison for tissue validation Region #1 and Region #2 (Supplementary Figure S9G) confirms the consistent agreement between U-Net prediction and SuRVoS segmentation.

### D. CT-based stereology

Following synchrotron imaging of a placental block, capturing 2520×2520×2120 px^3^ tissue (approximately 8 mm^3^) at ≈80 μm slice intervals with an image resolution of 0.8125 μm/px, systematic analysis of villous volume density and syncytiotrophoblast surface density was performed using a traditional stereology method, as described previously [58, 59]. Every 100^th^ image taken was imported into Image J software [60], providing 22 images in total (Figure S8A).

Within each image, a smaller 1800×1800 px^2^ field of view (FOV) was generated systematically (Figure S8B). A grid of 11×11 dots was superimposed on the FOV. The number of grid points hitting the villi and IVS was scored and the volume density of each of these morphological features was assessed using

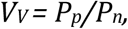

where *V*_*V*_ is the volume fraction of the feature of interest, *P*_p_ is the number of points that hit features of interest, and *P*_n_ is the total number of test points in the grid [59].

From each synchrotron image, a smaller 543×543 px^2^ field of view (FOV) was generated systematically (Figure S8C). A grid of 10×10 crosses (line length: 12 μm) was superimposed onto the FOV. In processing for surface density estimation, line intersects with the syncytiotrophoblast were scored (N = 100 intersects per image, N = 22 images). The horizontal lines were used as intercepts to estimate the specific surface area of syncytiotrophoblast *S*_*V (syn)*_ within tested volume:

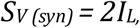

where *I*_*L*_ is the intersection count fraction (the number of intersections with the villous boundary per unit length of test line) [61].

### E. 3D morphological analysis

The structure of the fetal vascular network (Specimen 1) was analysed using Avizo software 2020.1 (Thermo Fisher Scientific). Using the *‘Centreline tree’* module which extracts the center lines of labelled 3D image volume as described in [62], the vascular network was skeletonized from placental tissue from the entire dataset of ≈8 mm^3^ (Figure 2A) and a small cropped region of ≈0.2 mm^3^ (Figure 2B). The vascular branching structure (length, diameter and tortuosity) of the entire network and a single connected tree inside the small cropped region was analysed. Only vessels with a diameter ≥3 px (2.4 μm) were included since the vessels smaller than this could not be resolved.

To evaluate the size and connectivity of the porous region (maternal IVS), the *‘Separate objects’* module (Avizo 2020.1) using a watershed method [63, 64] was employed on the labelled 3D image volume to separate the porous regions (Figure 2F). Afterwards, the *‘Pore Network Model’* module (Avizo) [62-64] was applied to generate the spheres at separated porous regions. The centre of two spheres were connected by ‘throat’ (a throat is a region that connects two individual pores) (Figure 2G).

In analysing the maternal porous region, the central placental tissue region of ≈1.8 mm^3^ (see supplementary Figure S3A) and the porous region that encompasses the single connected tree (from Figure 2C) were used. Distributions of diameters of pores and throat, length of throat and number of pore coordination (how many pores are connected to an individual pore) were analysed to characterize the placental porous medium.

The tortuosity of porous regions was analysed using the *‘Flow tortuosity’* module (Avizo) which computes the tortuosity based on the flow velocity vector field (the output of the *‘Absolute Permeability Experiment Simulation’* module [62, 63]). The flow velocity vector was calculated using a fixed pressure drop across the domain (see below for more details). Porous tortuosity was computed with various minimal lengths and compared between Specimen 1 and 2.

Maternal blood flow into the IVS was simulated using the *‘Absolute permeability experiment simulation’* module (Avizo 2020.1) which solves Stokes flow in a segmented intervillous porous space geometry [62, 63]. The simulation was performed on the small cropped region of ≈0.2 mm^3^ (as described above). However, the dataset was downsampled (2x) as flow simulation on full resolution data would be computationally prohibitive. The simulations were done in three directions (x, y and z) across the tissue thickness of ≈500 μm (x and z directions) and ≈950 μm (y direction). Three simulations were performed with a fixed pressure gradient across the three principal directions (pressure drop of 10 Pa in the y, or 5 Pa in the x and z directions respectively, accounting for ≈1:2:1 (x:y:z) aspect ratio; i.e. ≈10 Pa/mm within tissue [28, 29]). The other four domain boundaries (in the transverse to the applied gradient direction) were assumed impermeable in each simulation, and a no-slip flow condition was used at the fluid-solid interfaces. A constant blood viscosity of 0.003 Pa.s was assumed. The calculated net flow rate was used to estimate the empirical hydraulic permeability in each principal direction from Darcy’s law [62].

The area and volume fraction of segmented fetal tissue components from Specimen 1 and 2 were analysed using the *‘Volume fraction’* module (Avizo 2020.1). The surface area of segmented fetal tissue components was analysed using the *‘Label analysis’* module (Avizo) which calculates the surface area of labelled regions from 3D image volume. The area fraction was analysed for individual slices. Volume fraction and surface area were analysed using different region of interest (ROI) sizes. To obtain the ROI cubes at different sizes, the tissue cubes with various x and y length and fixed z thickness (243.75 μm) were cropped without overlapping. For those ROI sizes with more than 1000 ROI cubes, a systematic random sampling method (picking n^th^ ROI cubes) was used in order to get 1000 random ROI cubes for statistical analysis. For those ROI sizes with fewer than 1000 cubes, all ROI cubes were included. The analysis was done for both Specimen 1 and 2 and performed on the central tissue region (as shown in supplementary Figure S3A).

#### Statistical Handling

For the purposes of stereology, a systematic random sampling protocol was used, where every 100th image from a stack of 2120 images was analysed for volume fraction and specific surface area (N=22 fields of view).

## Acknowledgments

Imaging was performed on the Branchline I13-2 of the Diamond Light Source synchrotron in Oxfordshire, UK, on beamtimes MG23941 and MG22562. The authors would like to thank beamline scientists Dr Andrew Bodey and Dr Shashi Marathe and collaborators Dr Saurabh Shah and Prof Peter D. Lee (University College London) for providing access to the cold-stage and assistance with setting up. We also thank Dr James Carr, Dr Tristan Lowe, Dr Kerstin Schirrmann, Ms Ruth Whelan-Jeans and Ms Saskia Port (University of Manchester) for their help with pre-synchrotron optimisation of micro-CT protocols and image analysis.

## Author Contributions

WMT performed 3D image segmentation, data analysis and simulations, created movies and wrote the manuscript.

GP contributed to project design, data collection and data analysis, proposed optimised pipeline for specimen preparation and data segmentation and wrote the manuscript.

GN and HB refined and performed the perfusion and specimen collection, processed tissue and prepared blocks for imaging, and performed stereological analysis.

ONFK developed the U-Net segmentation algorithm and wrote the manuscript.

MB supervised development of U-Net segmentation algorithm, data segmentation, data analysis and movie creation.

YT developed and manufactured Yasuaki’s resin.

RML and EDJ contributed to the interpretation of results and provided critical feedback to shape the research, and wrote the manuscript.

PB conceived and designed the project, performed tissue sample collections, sample preparations and stereological analysis, contributed to the interpretation of the results, provided critical feedback, supervised the project and wrote the manuscript.

MCD supervised image segmentation, data analysis and simulations, contributed to the interpretation of the results, provided critical feedback, supervised the project and wrote the manuscript.

ILC conceived and designed the project, designed and performed the scale dependence error analysis, contributed to the interpretation of the results, supervised the project and wrote the manuscript.

All authors discussed the results and contributed to the writing of the manuscript.

## Data availability

All data needed to evaluate the results and conclusions are present in the paper and/or the Supplementary Materials. The reconstructed micro-CT volumes (DLS ID 120077 and 13761), manual segmentations (DLS ID 120077) and predicted final segmentations (DLS ID 123861) are available on public repository EMPIAR (https://www.ebi.ac.uk/pdbe/emdb/empiar/entry/10562/ and https://www.ebi.ac.uk/pdbe/emdb/empiar/entry/10563). The training data for U-net and trained U-Net models for segmenting the maternal/fetal blood volumes and the blood vessels are openly available at http://doi.org/10.5281/zenodo.4249627. The associated computational codes can be accessed at https://github.com/DiamondLightSource/python-placental-imaging. Additional data related to this paper may be requested from the authors.

## Ethics

All human placental tissues used in this study were acquired from term placentas delivered at St Mary’s Hospital, Manchester, with appropriate informed written consent and ethical approval (Tommy’s Project REC 15/NW/0829).

