## Supplementary data and text for "A massively multi-scale approach to characterising tissue architecture by synchrotron micro-CT applied to the human placenta"

#### SUPPLEMENTARY MATERIALS

##### S1. SAMPLE PREPARATION

**TABLE S1: Demographics and placental tissue preparation summary.**

| | Specimen 1 | Specimen 2 | Specimen 3 | Specimen 4 |
|---|---|---|---|---|
| DLS ID | 123761 | 120077 | 123744 | 123749 |
| Ethics (TOM) number | 2654-H2 | 2797-F | 2811-F | 3079-D |
| Mode of delivery [a] | CS | V | CS | CS |
| Maternal age (years) | 36 | 32 | 34 | 27 |
| Parity | 1 | 2 | 0 | 1 |
| Gestational age (weeks+days) | 39+0 | 37+6 | 35+2 | 39 |
| BMI | 24 | 29 | 25 | 30 |
| Smoker | No | No | Yes | No |
| Fetal sex | Male | Male | Male | Male |
| Birthweight (g) | 3386 | 2260 | 2082 | 3230 |
| Birthweight centile (%) | 49.2 | 0.3 | 4.6 | 29.9 |
| *Ex vivo* parameters | | | | |
| Cotyledon weight (g) | 29 | 39 | 25 | 40 |
| Fixative reagent | Z7+PFA | Z7+PFA | Z7+PFA | Z7+PFA |
| Contrast reagent – fetal side | Batson's | Yasuaki's | Yasuaki's | Yasuaki's |
| Contrast reagent – maternal side | Z7 | Z7 | Z7 | Z7+PTA |
| Tissue processing [b] | CP-dried | Frozen | Wax-embed | Wax-embed |

[a] CS = Cesarean; V = vaginal delivery
[b] Frozen = cryo-frozen in liquid N2 from fresh; CP-dried = ethanol dehydrated and critical-point dried; Wax-embed = ethanol dehydrated and wax-embedded

**A**

Segmentation routine proposed for the various methods of obtaining X-ray contrast

Biological structures of interest

| Structure | Segmentation routine | |
|---|---|---|
| Intervillous space | **Contrast based segmentation**<br>- Z7 fix & PTA stain | |
| | **Machine learning assisted segmentation**<br>- Z7 fix | |
| Stroma (syncytio-trophoblast) | **Contrast based segmentation**<br>- Extended Z7 fix or PTA stain | |
| | **Machine learning assisted segmentation**<br>- Extended Z7 fix or PTA stain | |
| Fetal villi network | **Contrast based segmentation** | |
| | **Vessels**: Vascular corrosion casts or perfuse with contrast agent containing resins. | **Capillary**: Perfuse with a low viscous resin to avoid artificially bulging out and corrosion cast or add nano-particles to resin for contrast. |
| | **Machine learning assisted segmentation** | |
| | **Vessels** | **Capillary** |
| | Resin perfuse and embed tissue (tissue corrosion or incorporation of contrast agent not necessary). | |

**Various methods for microCT placental sample preparation**

| | | |
|---|---|---|
| Specimen 1 - Ethanol dehydrate & critical point dry: May lead to tissue shrinkage. Sample motion from X-ray beam induced dehydration can be a potential issue when synchrotron source radiation is used. Gives the best contrast for syncytiotrophoblast. | Specimens 3 & 4 - Ethanol dehydrate & wax embed: May lead to tissue shrinkage and delamination of syncytiotrophoblast. Poor contrast between soft tissue and wax, requires phase contrast imaging. | Specimen 2 - Plunge freeze: Preserves physiological structures and is suited to imaging soft tissue. However, requires a cryo-stream or cold-stage to maintain samples under cryo-conditions during scanning. |

**B**

**Recommended pipeline for obtaining all the mesoscale placental features**

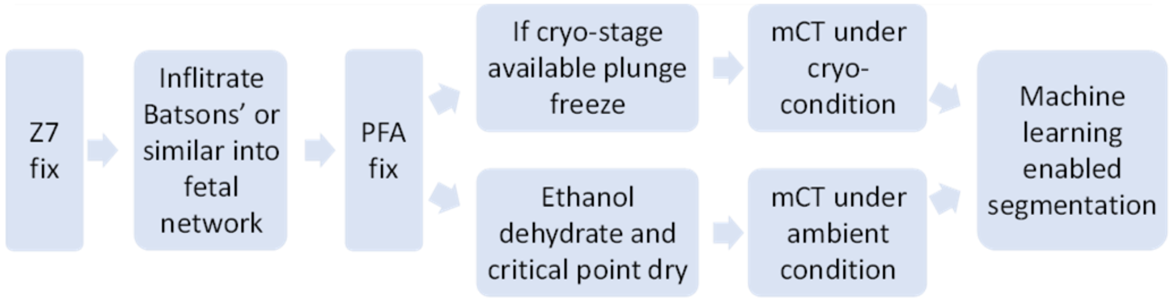


**FIG. S1:** Placental sample preparation required to confidently segment various features of the placenta. (A) Biological structures of interest and sample processing methods tried in this study. (B) Recommended sample processing pipelines to obtain various mesoscale placental features.

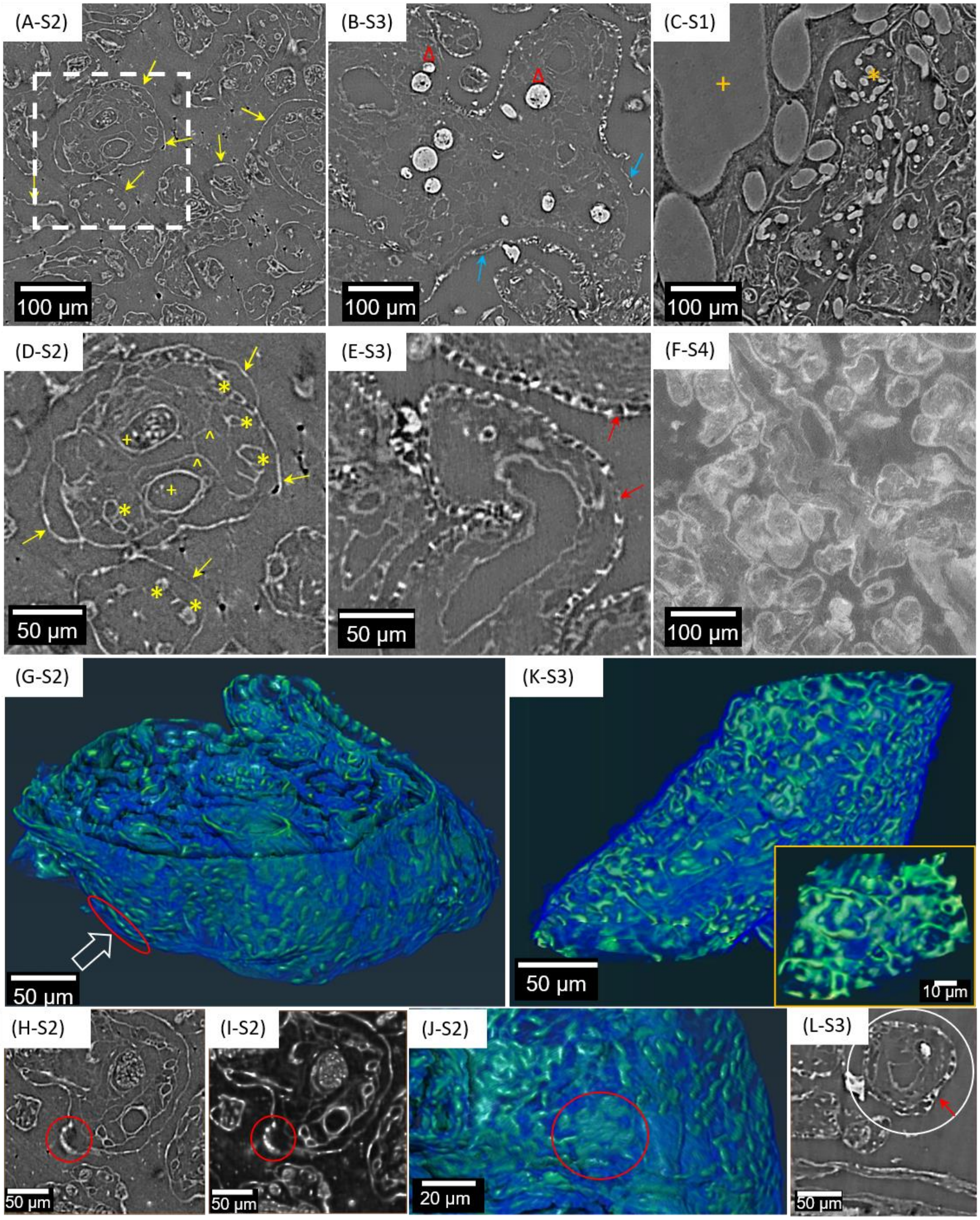


**FIG. S2: Two-dimensional cross-sectional images obtained from microCT data highlighting the influence of specimen preparation on the features that can be resolved.** The four specimens (labelled S1-S4; see Table S1) were imaged under three different conditions: plunge frozen in liquid nitrogen (A,D,G-J), ethanol dehydrated and wax embedded (B,E,F,K-L) and ethanol dehydrated and critical-point dried. (C). Plunge frozen specimens were imaged using a cold stage. All specimens were Zinc-7 fixed via maternal-side perfusion. (D) Magnified image of plunge frozen specimen (white box) shows the different placental architectures of a mature intermediate villous, including a well-resolved syncytiotrophoblast (arrows in D and 3D image in G), blood vessels (+; pair of arteriole with venule where one still contains red blood cells) and various capillaries (*). It is also possible to observe a network of the stroma (^). (G,J) 3D images of plunge frozen specimen showing the thin (~3 μm) syncytiotrophoblast envelope around the villous. (H,I) 2D cross-sectional images showing aggregated nuclei within a syncytial knot (red circle and on J showing 3D image in direction indicated by a white arrow in G). Ethanol dehydrated and either wax embedded (B,E,F) or critical-point dried (C) samples were imaged at ambient temperature. Wax embedded (B) and magnified images (E,L) show blood vessels, stroma and separately resolved microvillous and syncytiotrophoblasts (red arrows). (K) 3D images of the ethanol dehydrated and wax-embedded specimen showing an intermediate villous circled in (L). Inset of (K) showing the presence of pores/open channels on the syncytiotrophoblast envelope. Further, some of the blood vessels in (B) have been perfused with the Yasuaki resin (Δ). Blue arrows on (B) show part of the syncytiotrophoblast that has delaminated from the villous. Batson's resin, which lacks an X-ray contrast agent, can be observed in (C) with better perfusion of the vessels (+) and capillaries (*). Specimen 4 (F) produces the highest X-ray contrast for stroma and trophoblast after staining with phosphotungstic acid (PTA) for 120 hours, but with poor resolution of vascular features.

#### S2. IMAGE SEGMENTATION

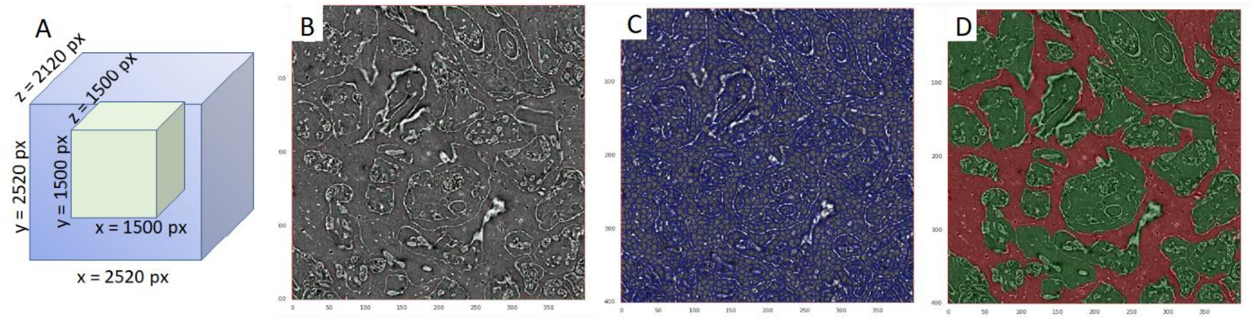


**FIG. S3: Fetal tissue segmentation and validation.** (**A**) The imaged tissue volume is ≈ 8 mm$^3$ (blue). Segmentations are of the central cropped region, with tissue volume of ≈ 1.8 mm$^3$ (green). (**B**) Example slice from original X-ray tomogram volume (Specimen 2). (**C**) Supervoxel regions creation based on filtered image features using SuRVoS. (**D**) Segmentation of fetal tissue and maternal IVS.

#### S3. FETAL VASCULAR ANALYSIS

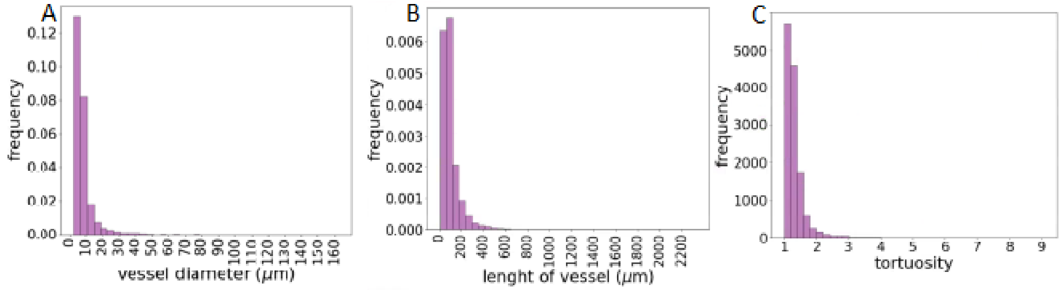


**FIG. S4: Additional characterisation of fetal vascular network.** Distribution of vessel (**A**) diameter, (**B**) length and (**C**) tortuosity of blood vessels from the entire vascular network from Specimen 1.

#### S4. MATERNAL POROUS REGION ANALYSIS

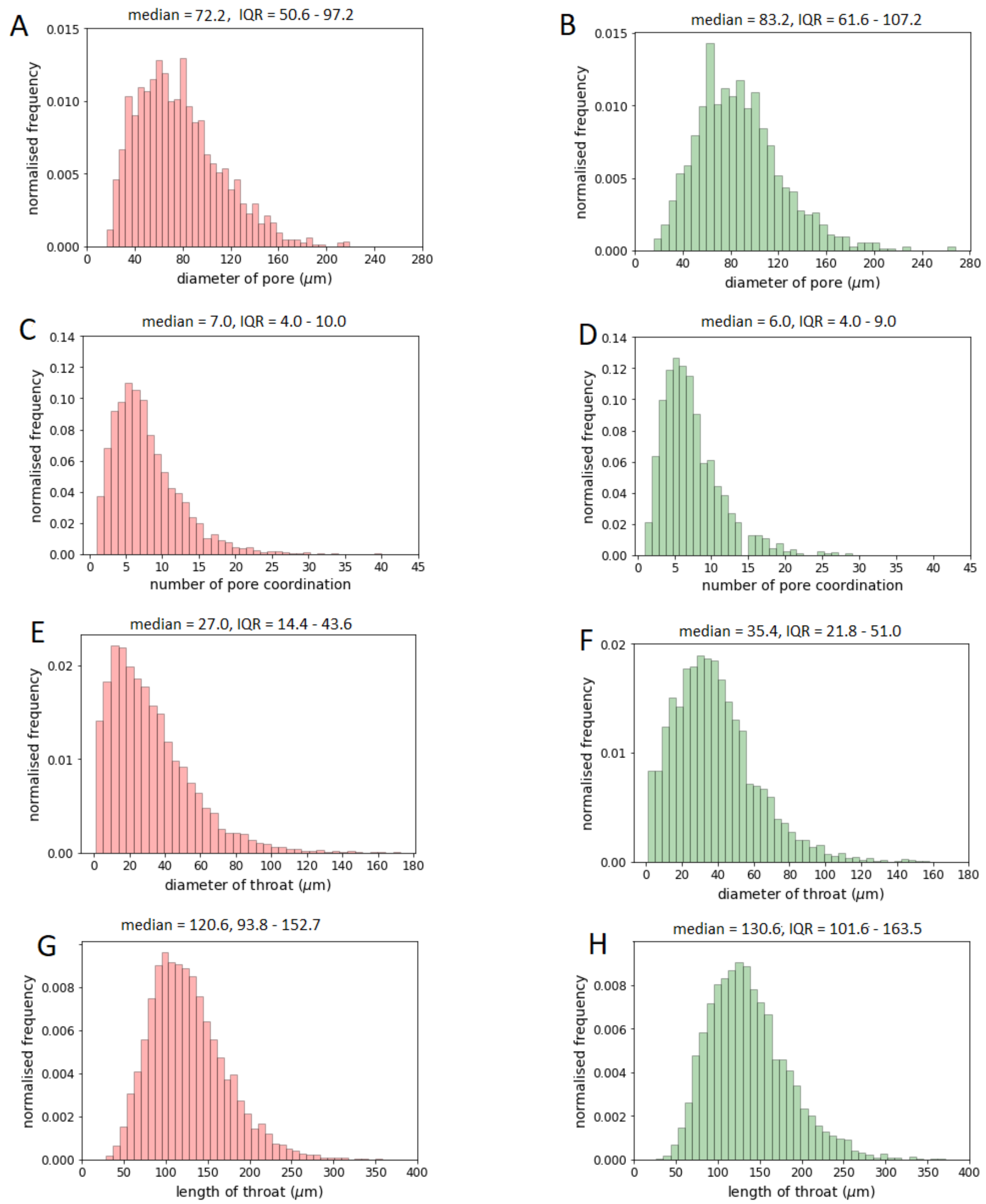


**FIG. S5: Additional characterisation of maternal porous regions.** Left column is specimen 1, right column is specimen 2. Distribution of (**A & B**) diameter of pores, (**C & D**) number of pore coordination, (**E & F**) diameter of throats and (**G & H**) length of throats. The median and interquartile range (in micrometer) are described on top of each graph.

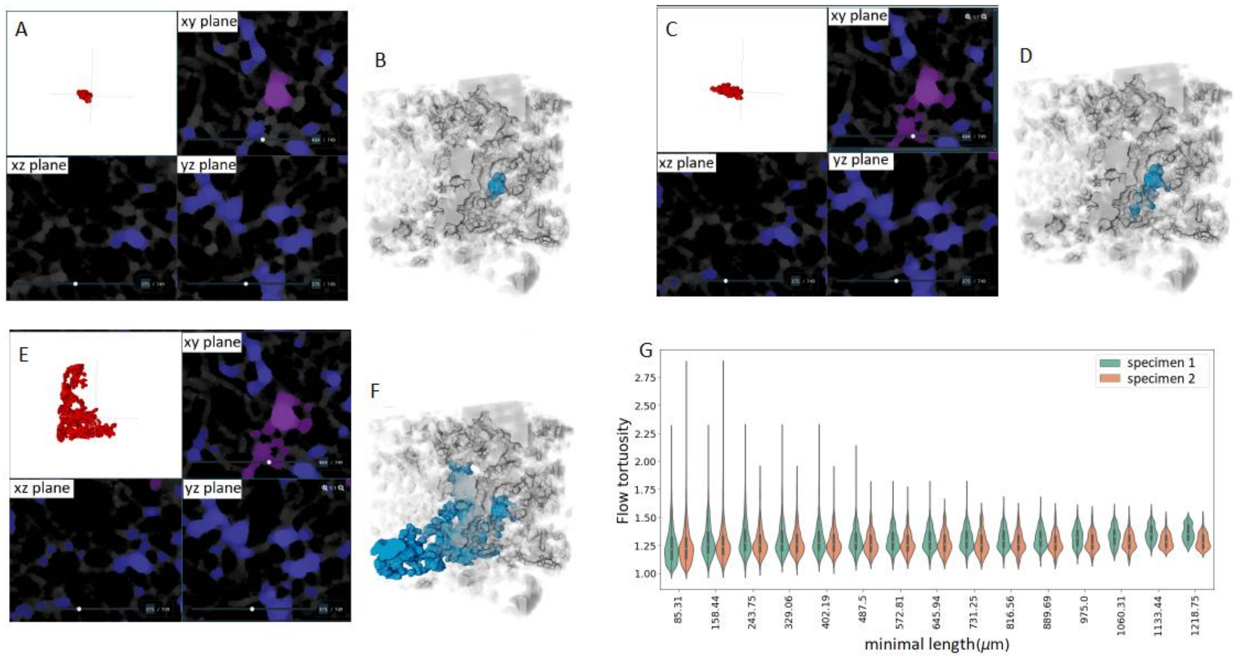


**FIG. S6: Assessment of pore connectivity (A-F) and tortuosity (G).** Using Specimen 2, a single pore is chosen in the tissue volume and its connectivity to other pores is limited by the diameter of the pore between (**A & B**) ≈ 48-110μm, (**C & D**) ≈ 40-110μm and (**E & F**) ≈ 37-110μm. a, c, d show the porous regions in three different planes and b, d and f show the association of porous regions with the entire volume. (**G**) Porous tortuosity (tortuosity of streamlines across the maternal IVS) measuring at various minimal lengths (of streamlines)**.**

#### S5. FETAL TISSUE ANALYSIS

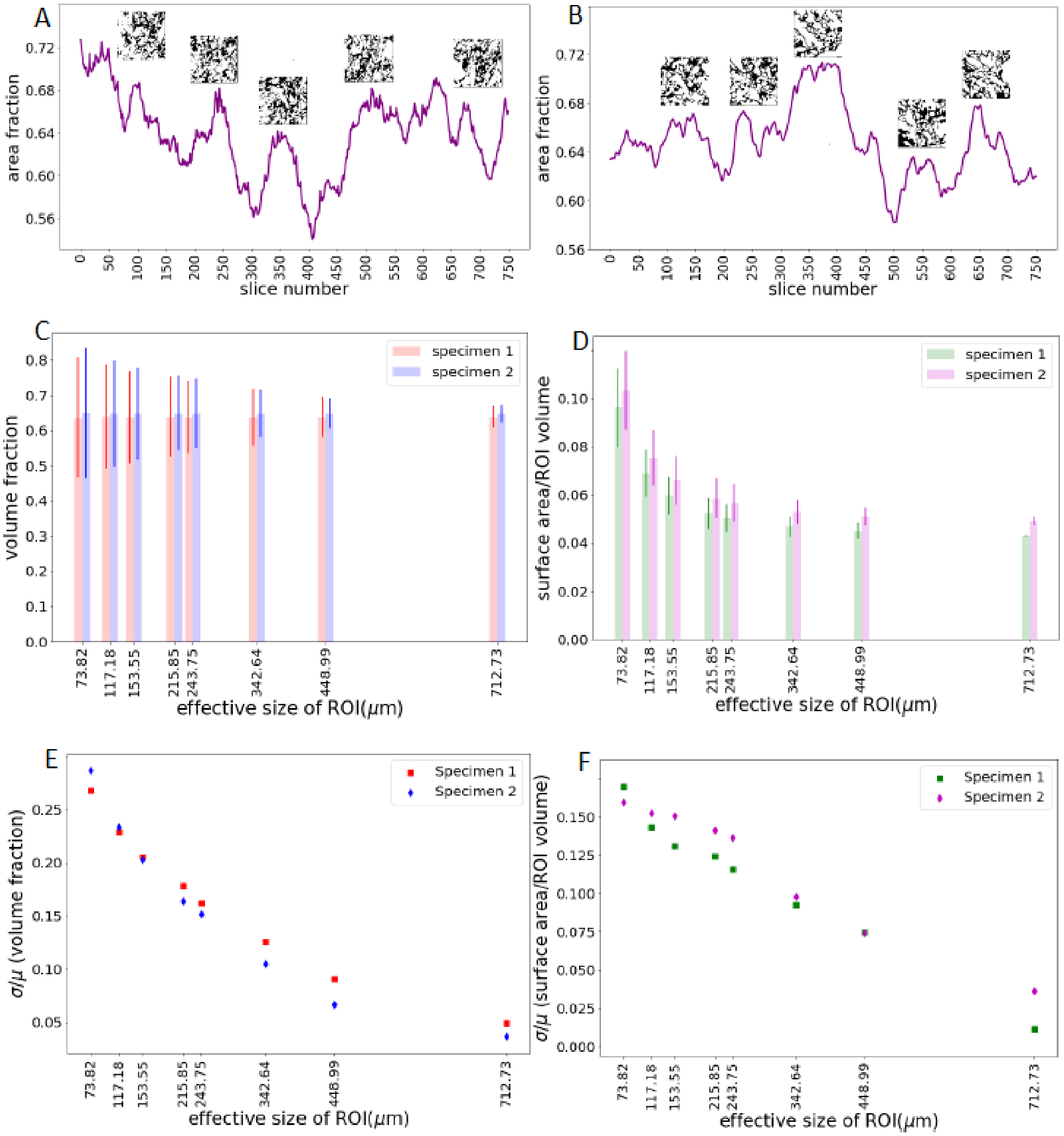


**FIG. S7: Additional characterisation of fetal tissue components.** (**A & B**) Area fraction variations in specimen 1 and 2. (**C**) Volume fraction vs. effective size of ROI (mean and std). (**D**) Ratio of surface area to ROI volume vs. effective size of ROI (mean and std). (**E & F**) Coefficient of variance of volume fraction and surface area/ROI volume.

#### S6. STEREOLOGICAL STUDY

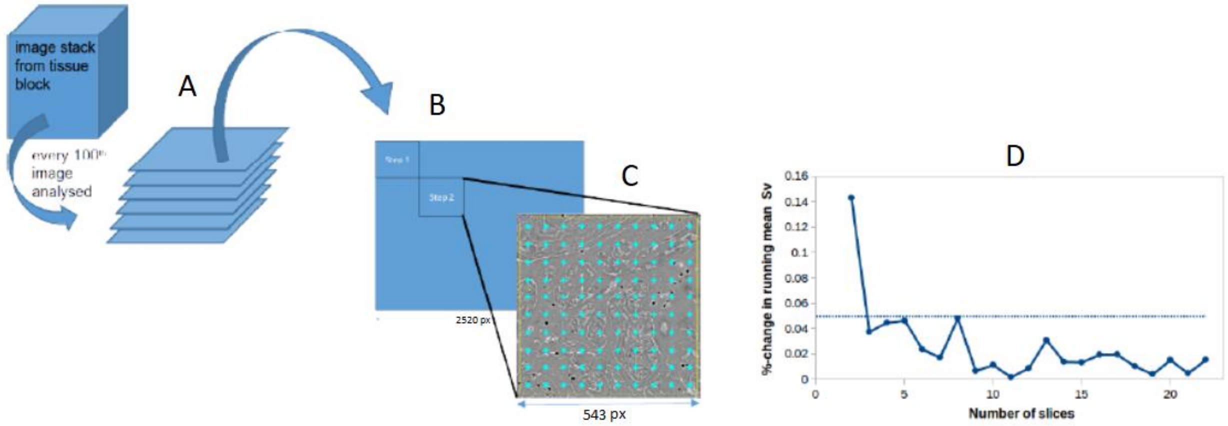


**FIG. S8: Systematic sampling strategy for stereology.** (**A**) Depicts the acquisition of images for analysis; (**B**) illustrates point analysis to yield volume density estimates of villi and IVS; and (**C**) shows line intersect processing to derive the surface density of the villi. Images are of a cryo-frozen human placental sample from a term FGR pregnancy. (**D**) Estimator convergence for the specific surface area (SV) of the villous tissue (analysed by stereological method from Specimen 2) with the number of slices; the dashed line denotes the 5% error level (relative change in the running mean).

#### S7. VALIDATION OF U-NET SEGMENTATION

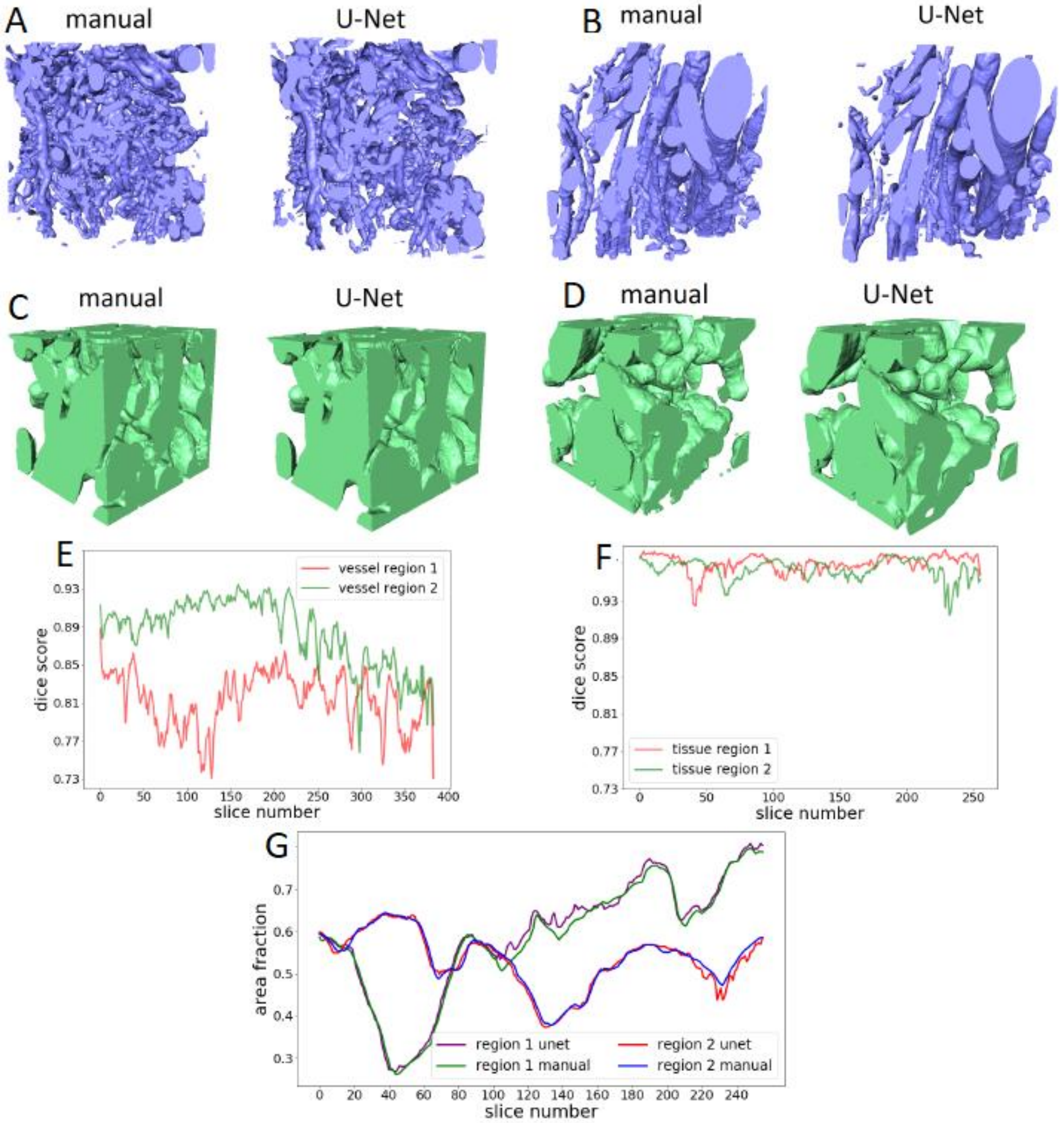


**FIG. S9: Validation of U-Net segmentation of fetal vascular network and fetal tissue components.** (**A & B**) Segmented volumes of vascular Region 1 and Region 2. (**C & D**) Segmented volumes of tissue Region 1 and Region 2. (**E**) Dice scores for vascular validation regions. (**F**) Dice scores for tissue validation regions. (**G**) Area fraction comparison between U-Net predictions and manual segmentations for tissue Region 1 and Region 2. All from Specimen 1.

### SUPPLEMENTARY METHODS

#### S8. CHARACTERISTICS OF TRANSPORT IN THE INTERVILLOUS SPACE.

Let us consider a region of interest of size $L \sim 1$ mm. For typical mobile solutes such as respiratory gases and small molecules, with diffusivity $D \sim 10^{-9}\,\mathrm{m^2/s}$ [65], assuming the mean intervillous space blood flow velocity $U$ of approximately $10\,\mu$m/s $\sim 10^{-5}\,\mathrm{m/s}$ (see Fig. 3), we have the Péclet number $Pe = U\,L/D \sim 10$, i.e. the flow contribution dominates molecular diffusion in the transport of solutes. Therefore, the pore structure, which determines the flow pathways in the intervillous space (see Figs 3 and S6), is physiologically important even for the most mobile solutes.

#### S9. UNCERTAINTY QUANTIFICATION OF STRUCTURAL METRICS IN A DISORDERED POROUS MEDIUM.

To characterise the scale-dependence of fluctuations in the structural metrics of a porous medium, consider a local volume fraction estimator $\hat{\phi}$ [43]

$$\hat{\phi}(\mathbf{x}) = \frac{1}{V_0} \int_{\mathscr{V}} I(\mathbf{z})\,\theta(\mathbf{z}-\mathbf{x})\;\mathrm{d}\mathbf{z}\,, \tag{S1}$$

where $I(\mathbf{x})$ is the indicator function for the phase of interest (say, the placental villous tissue; i.e. $I(\mathbf{x}) = 1$ for $\mathbf{x}$ in this phase, and is zero otherwise), and $\theta(\mathbf{x}) = \{1,\ \mathbf{x} \in \mathscr{V}_0;\ \ 0,\ \text{otherwise}\}$ is the indicator function for the observation region of interest (ROI) $\mathscr{V}_0$ of volume $V_0$, which is centred at an arbitrary location $\mathbf{x}$ within a larger porous medium domain $\mathscr{V}$.

For an ergodic and isotropic medium, $\hat{\phi}$ is shown to be an unbiased estimator, with $\mathbb{E}[\hat{\phi}] = \phi$, and the square root of normalized variance (termed ‘coarseness’) is given by [43]

$$\frac{\sigma_{\hat{\phi}}}{\phi} = \frac{1}{\phi\,V_0}\left[\iint_{\mathscr{V}} \left[S_2(\mathbf{r}) - \phi^2\right]\;V_{\mathrm{int}}(\mathbf{r})\;\mathrm{d}\mathbf{r}\right]^{1/2}\,, \tag{S2}$$

where $S_2(\mathbf{r}) = \mathbb{E}[I(\mathbf{x})I(\mathbf{x}+\mathbf{r})]$ is the two-point probability function, which approaches the value of $\phi^2$ for sufficiently large $r$, and $V_{\mathrm{int}}(\mathbf{r})$ is the intersection volume of two ROIs separated by the distance $r$, which depends both on the ROI size and on its shape [43].

Following Torquato *et al.* [43, 66], in the limit of sufficiently large ROI volumes (i.e. when the ROI size $L$ is much larger than the intrinsic correlation lengthscale $\lambda$, such that $S_2(\lambda) \approx \phi^2$), the long-range structural correlations become negligibly small, and the integral (S2) reduces to an integral in the vicinity of $r \leq \lambda$. In this case, the intersection volume $V_{\text{int}} \approx V_0\,(1 - \mathcal{O}(\lambda/L))$ approaches the volume $V_0$ of the ROI, and (S2) gives an asymptotic relationship

$$\frac{\sigma_{\hat{\phi}}}{\phi} \sim {V_0}^{-1/2}\, f(\phi, \lambda)\,. \tag{S3}$$

Similarly, we introduce an estimator $\hat{s}$ for the specific surface area $s$ of the tissue phase:

$$\hat{s}(\mathbf{x}) = \frac{1}{V_0} \int_{\mathcal{V}} |\nabla I(\mathbf{z})|\, \theta(\mathbf{z} - \mathbf{x})\; \mathrm{d}\mathbf{z}\,, \tag{S4}$$

where $|\nabla I(\mathbf{x})|$ is the characteristic function for the interface, defined in the sense of generalised functions [66, 67], so that $\mathbb{E}[|\nabla I(\mathbf{x})|] = s$. Utilising the surface-surface probability function $F_{ss}(\mathbf{x}) = \mathbb{E}[|\nabla I(\mathbf{x})|\, |\nabla I(\mathbf{x}+\mathbf{r})|]$ ($F_{ss} \to s^2$ as $r \to \infty$) [67], and assuming an ergodic statistically homogeneous porous medium, we extend the technique of Lu and Torquato [43] to derive an analogous to (S2) relative variability for the specific surface area:

$$\frac{\sigma_{\hat{s}}}{s} = \frac{1}{s\,V_0} \left[ \iint_{\mathcal{V}} \left[F_{ss}(\mathbf{r}) - s^2\right]\, V_{\text{int}}(\mathbf{r})\; \mathrm{d}\mathbf{r} \right]^{1/2}, \tag{S5}$$

where $\sigma_{\hat{s}}^2 \equiv \mathbb{V}\mathrm{ar}[\hat{s}]$.

By taking the limit of large ROI volumes for an isotropic and homogeneous porous medium, i.e. for the ROI size $L \gg \lambda_s$, we have $F_{ss}(\lambda_s) - s^2 \approx 0$ and $V_{\text{int}} \approx V_0$, and thus the relationship (S5) can be approximated by

$$\frac{\sigma_{\hat{s}}}{s} \sim {V_0}^{-1/2}\, g(s, \lambda_s)\,, \tag{S6}$$

where, similarly to (S3), the relative magnitude of the fluctuations drops as ${V_0}^{-1/2}$, with a factor that only depends on the correlation lengthscale and the expectation of the specific surface area.
